# scSpatialSIM: a simulator of spatial single-cell molecular data

**DOI:** 10.1101/2024.02.08.579486

**Authors:** Alex C Soupir, Julia Wrobel, Jordan H. Creed, Oscar E Ospina, Christopher M Wilson, Brandon J Manley, Lauren C. Peres, Brooke L Fridley

## Abstract

**Background:** Spatial molecular data is increasingly being generated in biological tissue studies to increase our understanding of cell infiltration and spatial architecture of tissues. Examples of technologies used to study the spatial contexture of tissues are single-cell protein expression assays and spatial transcriptomics. The increased use of spatial biology technologies has also resulted in an increase in the development of statistical methods to describe the spatial landscape in tissues. Due to the lack of consensus on “gold standard” statistical approaches for assessing the spatial contexture of tissues, we created an R package, *scSpatialSIM*, to assess different statistical and bioinformatic methods. *scSpatialSIM* allows users to simulate single-cell molecular data to mimic real tissues at scale, clustering of cell types, and co-clustering / co-localization of two or more cell types. *scSpatialSIM* also contains functions that give users the ability to simulate quantitative distributions for positive and negative cells (e.g., gene expression, fluorescence intensity).

**Results:** We demonstrate that *scSpatialSIM* allows users to easily simulate various kernel densities of probability distributions used to create the marked point pattern – points distributed in space with either numeric or categorical features. Using *scSpatialSIM*, we used four univariate spatial simulation scenarios to compare three different measures for spatial clustering (Ripley’s K(*r*), nearest neighbor G(*r*), and pair correlation g(*r*)). We found that Ripley’s K(*r*) identifies the most radii with significant clustering in all four scenarios. Nearest neighbor G(*r*) only identified all samples as significantly clustered at one radius (*r* = 0.07) in one simulation scenario (high abundance large cluster size). Pair correlation g(*r*) was better able to detect significant clustering at low radii when abundance was low.

**Conclusions:** Vignettes developed for *scSpatialSIM* cover the creation of single-type and multi-type spatial single-cell molecular data, as well as how these simulated data can be used with other R packages, such as *spatialTIME*, to derive spatial statistics. Development of this package is crucial for furthering our understanding of the power of existing methods and the development of novel applications to assess the spatial contexture of tissues by providing an objective platform for simulating spatial single-cell molecular data.

## Background

Studying the spatial contexture of cells in biological tissues is increasing in popularity among biomedical researchers. New technologies allow the profiling and phenotyping of cells in tissue microarrays (TMAs) and whole tissue slides to understand the spatial architecture of tissues, such as the tumor microenvironment (TME), and the distributions of cells in 2-diminsional space. The ability to characterize the spatial context of cells has allowed for further exploration of the correlation between the tissue architecture and patient outcomes, such as overall survival or response to therapies [1-3]. In contrast to studying only the cell abundance or density, the spatial contexture of cell types and their level of aggregation or dispersion can provide more details about the organization of complex tissues and answer important questions about cell-cell communication and their associations [3, 4].

Single-cell protein imaging platforms, such as multiplex immunofluorescence (mIF) or immunohistochemistry (mIHC) [5], have been successfully used to profile cell types in tissues, as well as spatially characterize the immune microenvironment of several malignancies [3, 6]. In addition to single-cell protein data, spatial resolved transcriptomics (SRT) has recently gained traction in biological research. SRT experiments can range from studies involving imaging or sequencing regions of tissue (GeoMx, [7], Nanostring, Seattle, WA) to equally spaced grids with unique molecular identifiers (UMIs) and spatial barcodes containing between 5 and 10 cells (Visium [8], 10x Genomics, Pleasanton, CA) to sub-cellular transcript locations (CosMx Spatial Molecular Imaging [9], Nanostring, Seattle, WA; Xenium [10], 10x Genomics). These spatial molecular technologies require complex statistical and bioinformatic methods to summarize the spatial characteristics of the tissues. Statistical methods from the field of ecology have been used to assess the level of clustering of cell types in tissues, but there is a lack of consensus on which method is the “gold standard”.

To bridge this gap in the literature, we have developed an R package, *scSpatialSIM*, that allows for fine-tuned simulation of spatial single-cell molecular data; these can be leveraged to compare available statistical methods for measuring single-cell spatial data. While packages like *spaSim* exist to simulate cell-level data to mimic tissue architectures, *scSpatialSIM* can also simulate continuous data (e.g., mIF intensity data, SRT gene expression data) depending on user input and parameter settings [11]. Our simulation framework conceptualizes cell spatial distributions as marked point patterns generated by an underlying point process that controls the spatial clustering and repulsion of cells and cell subtypes. Simulated mIF and SRT datasets generated by our package allow for benchmarking of statistical methods to determine the power of these approaches to detected different levels of clustering, co-localization, or segregation of cell types.

## Implementation

*scSpatialSIM* is an R package that is available for installation on R4.0.0 [12] or later from GitHub (https://github.com/fridleylab/scSpatialSIM) and CRAN. The functions within *scSpatialSIM* support “piping” (i.e., using the magrittr ‘%>%’ function), which allows function calls to be chained together to develop whole simulation pipelines from start to finish. This feature is possible with the implementation of the spatial simulation object (S4 class SpatSimObj). The SpatSimObj serves as storage for all the parameters and results performed with *scSpatialSIM*. Our goal was to provide a collection of methods that allow for the creation of realistic simulations of cell types within tissue samples in a stepwise manner: 1) point patterns, 2) tissue regions or domains, 3) holes/non-cellular areas, and 4) cell phenotypes (**Figure 1**). The SpatSimObj also stores the simulation space (class ‘owin’ from *spatstat* [13]), the number of independent simulations to perform, and the number of cell types to simulate. Additionally, we have created vignettes (https://fridleylab.github.io/scSpatialSIM) that demonstrate how to use *scSpatialSIM* as well as how the outputs of *scSpatialSIM* can be input into other R packages to derive spatial statistics, such as with the R package *spatialTIME* [14].

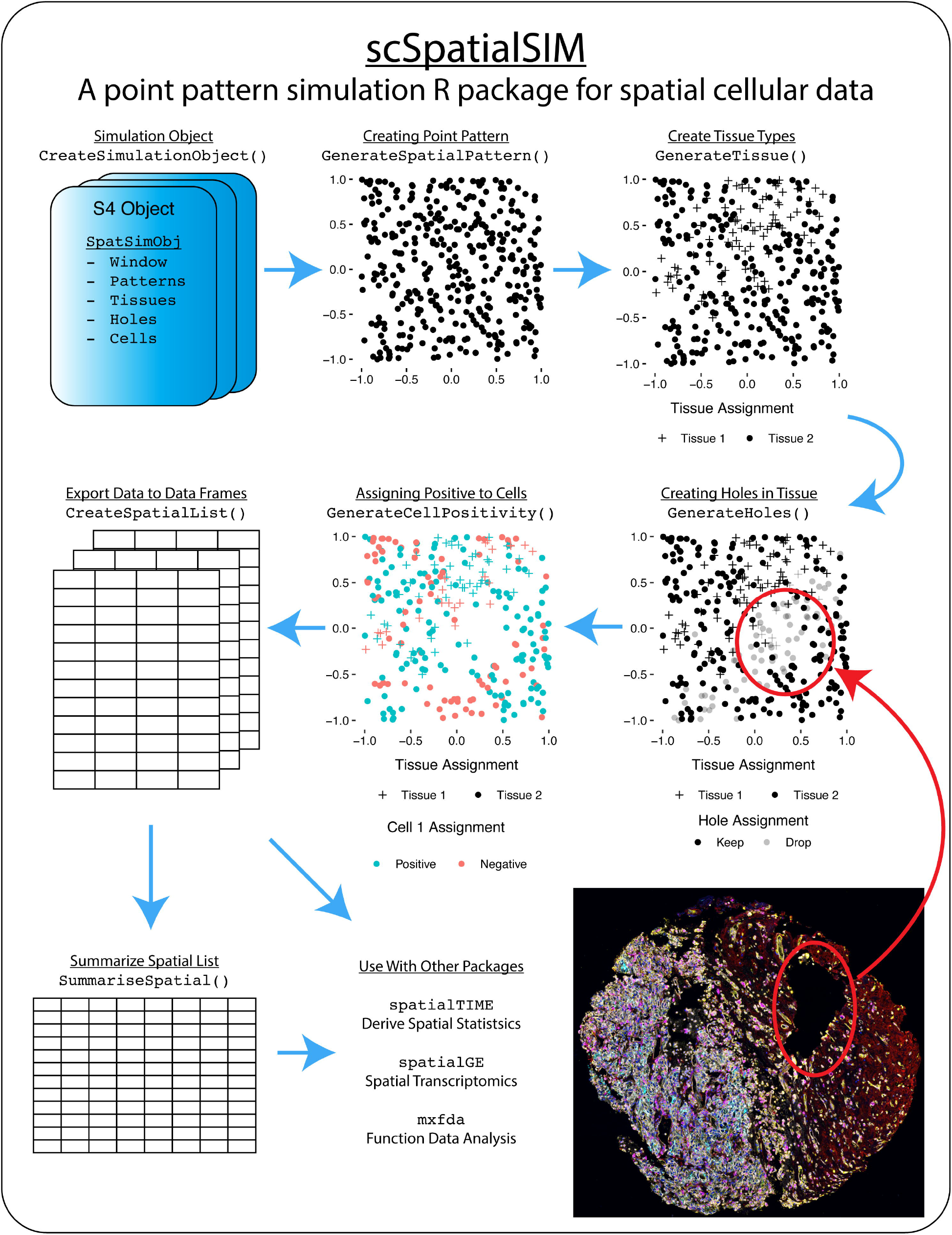

Following the creation of the spatial simulation object, the point pattern is created using ‘GeneratePointPattern()’ which calls the ‘rpoispp()’ function from *spatstat*. All other point classifications (i.e., creating a marked point process) are performed by randomly creating probability surface using Gaussian kernels within user provided constraints. If no parameter options are provided, single and multi-type simulations use the default parameters to set the kernels for the creation of the probability surface. Tissue class assignment allows for two different tissue types or domains (i.e., tumor vs stroma compartment) to be assigned on each point pattern. Similarly, the package allows for the generation of regions in which no cells are measured due to technical and biological reasons (e.g., folded regions of tissue, holes) which may be common in biological tissue samples constructed into tissue microarrays (TMAs) [3]. Differences between single- and multi-type cell simulations comes from the ability to adjust the level of colocalization or segregation of cell types. Using the function ‘GenerateCellPositivity()’, users provide a ‘shift’ parameter which allows for co-localization or spatial segregation of simulated cell types. This is a critical parameter allowing benchmarking of statistical methods to assess their ability to detect clustering or colocalization of different cell types.

We have also provided methods for exporting the data within the SpatSimObj in tabular form for use by other analysis software or R packages, such as the R package *spatialTIME* [14]. For example, traditionally there are spatial files that represent each sample in their entirety as well as a summary file that contains the number of total cells positive for markers and their corresponding proportion of total cells. *scSpatialSIM* can export these data from its S4 object into a summary data frame and list of sample level data frames. This makes it easier to simulate numerous scenarios of cell clustering in tissues and use external software to test analytical approaches. Additionally, different distributions can be generated by cell positivity (such as fluorescence intensity or gene expression) using the exported data with ‘GenerateDistributions()’. For example, fluorescence intensity of FOXP3 protein marker to determine regulatory T cell phenotype status in spatial single-cell protein studies.

## Results

Using *scSpatialSIM*, we created four simulation scenarios with either high or low kernel density (i.e., high or low abundance of the cell type of interest) combined with either high or low kernel variance (i.e., cells are in a tightly clustered pattern or cells are in a more dispersed clustered pattern) to represent tissues with different cell abundances and cluster sizes with 1000 simulated samples each. Using these simulations, we applied spatial summary functions within the *spatialTIME* R package (Ripley’s K(r), nearest neighbor G(r), and pair correlation g(r)) with 100 permutations to demonstrate the ability of the summary functions to identify significant spatial clustering of positive cells (over complete spatial randomness) at radii of 0 to 0.5 in increments of 0.01 (**Figure 2**). **Table 1** shows the number of radii for which each summary function had the highest frequency of significant samples, indicating that Ripley’s K(r) was able to identify cell-type clustering more often than the other two measures. We also found in simulations with high kernel density (high abundance of cells positive for a marker) that Ripley’s K(r) identified 100% of samples showing significant clustering across 24 and 46 radii for small and large clusters sizes (i.e., variance in the kernel), respectively. Conversely, at no radius did the nearest neighbor G(r) measure identify more samples with significant clustering than Ripley’s K(r) or pair correlation g(r), and only one radius (r=0.07) identified clustering in the high abundance/large cluster size simulation with 100% of samples significantly clustered.

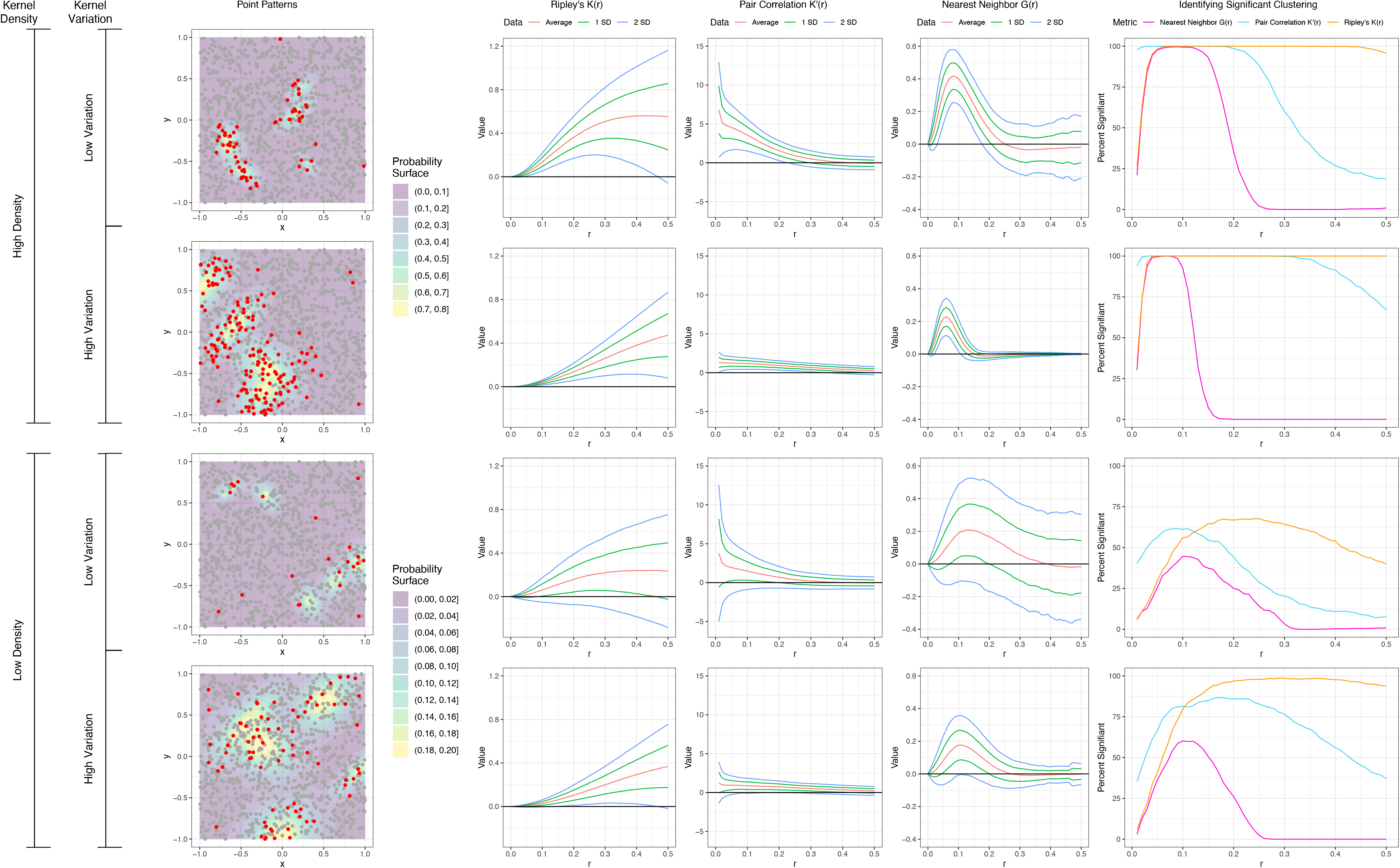

## Discussion

*scSpatialSIM* is an R package that provides users with a simplified coding interface to simulate spatial single-cell molecular data. It provides functions that allow adjustment of cell clustering, tissue assignment, and tissue holes, along with the ability to simulate continuous measurements. The package also allows for easy exporting of simulated data to be used downstream in other software packages. To illustrate the use of *scSpatialSIM*, we demonstrated its use in benchmarking univariate point pattern summary functions Ripley’s K(r), nearest neighbor G(r), and pair correlation g(r). *scSpatialSIM*’s intended use is to simulate spatial single-cell molecular data for benchmarking new statistical and bioinformatics approaches to summarize the spatial architecture of tissues.

## Conclusions

Simulation of realistic spatial single-cell molecular data allows for benchmarking of different spatial statistics in a biologically meaningful context. Increased use of technologies that provide biological information within a spatial context will undoubtably result in new analytical approaches to summarize spatial information primed for further scientific study to define their biological consequences. The R package *scSpatialSIM* provides a platform to initiate these comparisons at scale with the ability to fine tune different features of the spatial architecture in the simulated tissue sample.

## Declarations

### Ethics approval and consent to participate

Not applicable.

### Consent for publication

Not applicable.

### Availability of data and materials

Source code is available on GitHub (https://github.com/FridleyLab/scSpatialSIM) and the package can be installed in R with ‘install.packages()’ from the CRAN repository (https://cran.r-project.org/web/packages/scSpatialSIM).

### Authors’ contributions

BLF and CMW conceptualized the method of simulating spatial single-cell data. ACS, JHC, and CMW wrote the R code and designed the objects for the package. BJM and LP contributed data for realistic tissue for which to emulate with simulations. JW and OEO independently tested the package. ACS and BLF designed and performed the univariate spatial metric comparison. ACS and BLF drafted the manuscript and all authors contributed to edits and agreed on final version.

## Availability and requirements

Project name: *scSpatialSIM* a point pattern simulator for single-cell spatial molecular data Project home page: https://github.com/fridleylab/scSpatialSIM; https://cran.r-project.org/web/packages/scSpatialSIM/

Operating system(s): Windows, Linux, Mac

Programming language: R

Other requirements: R >= 4.00

License: MIT

Any restrictions to use by non-academics: None

